# Checkpoint inhibition of origin firing prevents inappropriate replication outside of S-phase

**DOI:** 10.1101/2020.09.30.320325

**Authors:** Mark C. Johnson, Geylani Can, Miguel Santos, Diana Alexander, Philip Zegerman

## Abstract

Across eukaryotes, checkpoints maintain the order of cell cycle events in the face of DNA damage or incomplete replication. Although a wide array of DNA lesions activates the checkpoint kinases, whether and how this response differs in different phases of the cell cycle remains poorly understood. The S-phase checkpoint for example results in the slowing of replication, which in the budding yeast *Saccharomyces cerevisiae* is caused by Rad53 kinase-dependent inhibition of the initiation factors Sld3 and Dbf4. Despite this, we show here that Rad53 phosphorylates both of these substrates throughout the cell cycle at the same sites as in S-phase, suggesting roles for this pathway beyond S-phase. Indeed we show that Rad53-dependent inhibition of Sld3 and Dbf4 limits re-replication in G2/M phase, preventing inappropriate gene amplification events. In addition we show that inhibition of Sld3 and Dbf4 after DNA damage in G1 phase prevents premature replication initiation at all origins at the G1/S transition. This study redefines the scope and specificity of the ‘S-phase checkpoint’ with implications for understanding the roles of this checkpoint in the majority of cancers that lack proper cell cycle controls.

## Introduction

It is vitally important that in every cell division the entire genome is replicated once and only once. In eukaryotes this is achieved by linking DNA replication control to the cell cycle (Siddiqui et al., 2013). The first step in replication is the formation of the pre-replicative complex (pre-RC) at origins – a process called ‘licensing’. Licensing involves the Orc1-6 and Cdc6-dependent loading of double hexamers of the Mcm2-7 helicase on double stranded DNA. Licensing is restricted to late mitosis/early G1 phase by the activity of the APC/C, which eliminates licensing inhibitors such as cyclin-dependent kinase (CDK) and geminin in this window of the cell cycle. In the budding yeast *Saccharomyces cerevisiae*, which lacks geminin, CDK inhibits licensing from late G1 phase until mitosis by multiple mechanisms including direct phosphorylation of Orc2/Orc6, nuclear exclusion of the Mcm2-7 complex and by mediating SCF^CDC4^-dependent degradation of Cdc6 (Blow and Dutta, 2005; Nguyen et al., 2001).

Importantly, Mcm2-7 double hexamers loaded in late M/early G1 phase are inactive and replication initiation can only occur after the inactivation of the APC/C at the G1/S transition. APC/C inactivation allows the accumulation of S-phase CDK and Dbf4-dependent (DDK) kinase activities (Labib, 2010). DDK directly phosphorylates the inactive Mcm2-7 double hexamers, generating a binding site for firing factors including Sld3/Sld7 and Cdc45, while CDK phosphorylates Sld3 and an additional initiation factor Sld2, which via phospho-interactions with Dpb11, results in replisome assembly by poorly understood mechanisms (Riera et al., 2017). This duality of function of CDK, both as an inhibitor of licensing and as an activator of the replisome is critical to ensure once per cell cycle replication (Diffley, 2004).

In light of the importance of the linkage between DNA replication control and cell cycle progression, multiple checkpoints exist to regulate DNA synthesis and genome integrity before (G1 checkpoint), during (S-phase checkpoint) and after S-phase (G2/M checkpoint, Hartwell and Weinert, 1989; Kastan and Bartek, 2004). These checkpoints are mediated by the PI3 kinase superfamily checkpoint kinases ATM/ATR (Tel1/Mec1 in budding yeast) and the effector checkpoint kinases Chk1/Chk2 (Chk1/Rad53 in budding yeast).

In G1 phase, DNA damage such as UV photoproducts causes checkpoint-dependent delays in the onset of DNA replication by inhibition of G1/S cyclins (Lanz et al., 2019; Shaltiel et al., 2015). In budding yeast this occurs in part by Rad53-dependent phosphorylation and inhibition of the Swi6 subunit of the transcriptional activator SBF (SCB binding factor) leading to reduced cyclin transcription (Sidorova and Breeden, 1997) and in humans by ATM-Chk2 mediated stabilisation of p53, as well as by checkpoint-dependent degradation of cyclin D and Cdc25A (Lanz et al., 2019; Shaltiel et al., 2015).

Although CDK and DDK are activated at the G1/S transition, normally origin firing occurs as a continuum throughout S-phase, with some origins firing in the first half of S-phase (early origins) and others in the second half (late origins). When replication forks emanating from early firing origins stall, for example due to DNA lesions, activation of the S-phase checkpoint kinase response results in the dramatic slowing of replication rates (Painter and Young, 1980; Paulovich and Hartwell, 1995), which occurs in large part through inhibition of late firing origins (Yekezare et al., 2013). In budding yeast, Rad53 blocks late origin firing by directly inhibiting two replication initiation factors; the DDK subunit Dbf4 and the CDK target Sld3 (Lopez-Mosqueda et al., 2010; Zegerman and Diffley, 2010). The checkpoint-mediated inhibition of origin firing likely occurs by similar mechanisms in human cells as the checkpoint kinases also bind to and inhibit the Sld3 orthologue Treslin (Boos et al., 2011; Guo et al., 2015) and inhibit DDK (Costanzo et al., 2003; Lee et al., 2012). One function of inhibiting origin firing during S-phase in the presence of DNA lesions is to prevent the exhaustion of essential factors, such as topoisomerase activities, by excessive numbers of replisomes (Morafraile et al., 2019; Toledo et al., 2017).

A key proposed feature of the DNA damage checkpoints is that the response is tailored to the cell cycle phase in which the DNA damage occurred. Despite this, there is very little evidence to suggest that substrate specificity of the checkpoint kinases changes during the cell cycle. Indeed, in budding yeast most forms of DNA damage and replication stress converge on the single effector kinase Rad53, but how different checkpoint responses in different cell cycle phases can be mediated by a single kinase is not known. In this study we set out to explore the specificity of Rad53 towards the replication substrates Sld3 and Dbf4 across the cell cycle in the budding yeast *Saccharomyces cerevisiae*. We show that Rad53 phosphorylates both of these substrates throughout the cell cycle at the same sites as in S-phase. From this we hypothesised that although these substrates are deemed to be targets of the ‘S-phase checkpoint’, Rad53 may also prevent aberrant origin firing outside of S-phase. Indeed we show that Rad53-dependent inhibition of Sld3 and Dbf4 limits re-initiation of replication in G2/M phase and also prevents premature firing of all origins, not just late origins, at the G1/S transition. This study overhauls our understanding of the cell cycle phase specificity of the ‘S-phase checkpoint’ and provides a novel mechanism that restricts replication initiation to a specific window of the cell cycle after DNA damage.

## Results and Discussion

### Sld3 and Dbf4 are phosphorylated by Rad53 outside of S-phase

Since the DNA damage checkpoint response can be activated in all phases of the cell cycle, we addressed whether the replication factors Sld3 and Dbf4 could be targeted by Rad53 outside of S-phase *in vivo* in budding yeast. To test this, we first analysed the consequences of DNA damage in G1 phase cells arrested with the mating pheromone alpha factor (Figure 1A). These experiments were conducted with strains containing a null mutation in the alpha factor protease, *bar1Δ*, to ensure that cells were fully arrested in G1 phase and had not started DNA replication. Addition of the UV mimetic drug 4-NQO to G1 phase cells resulted in robust Rad53 activation, as determined by the accumulation of the phospho-shifted forms of the kinase (Figure 1B, 1C). Importantly, we observed a dramatic increase in lower mobility forms of Sld3 when Rad53 was activated in G1 phase (Figure 1B), which was indeed Rad53-dependent (Supplementary Figure 1). For Dbf4, which is an APC/C substrate and partially degraded in alpha factor arrested cells (Ferreira et al., 2000), we also observed a mobility shift in G1 phase coincident with Rad53 activation (Figure 1C).

**Figure 1.**
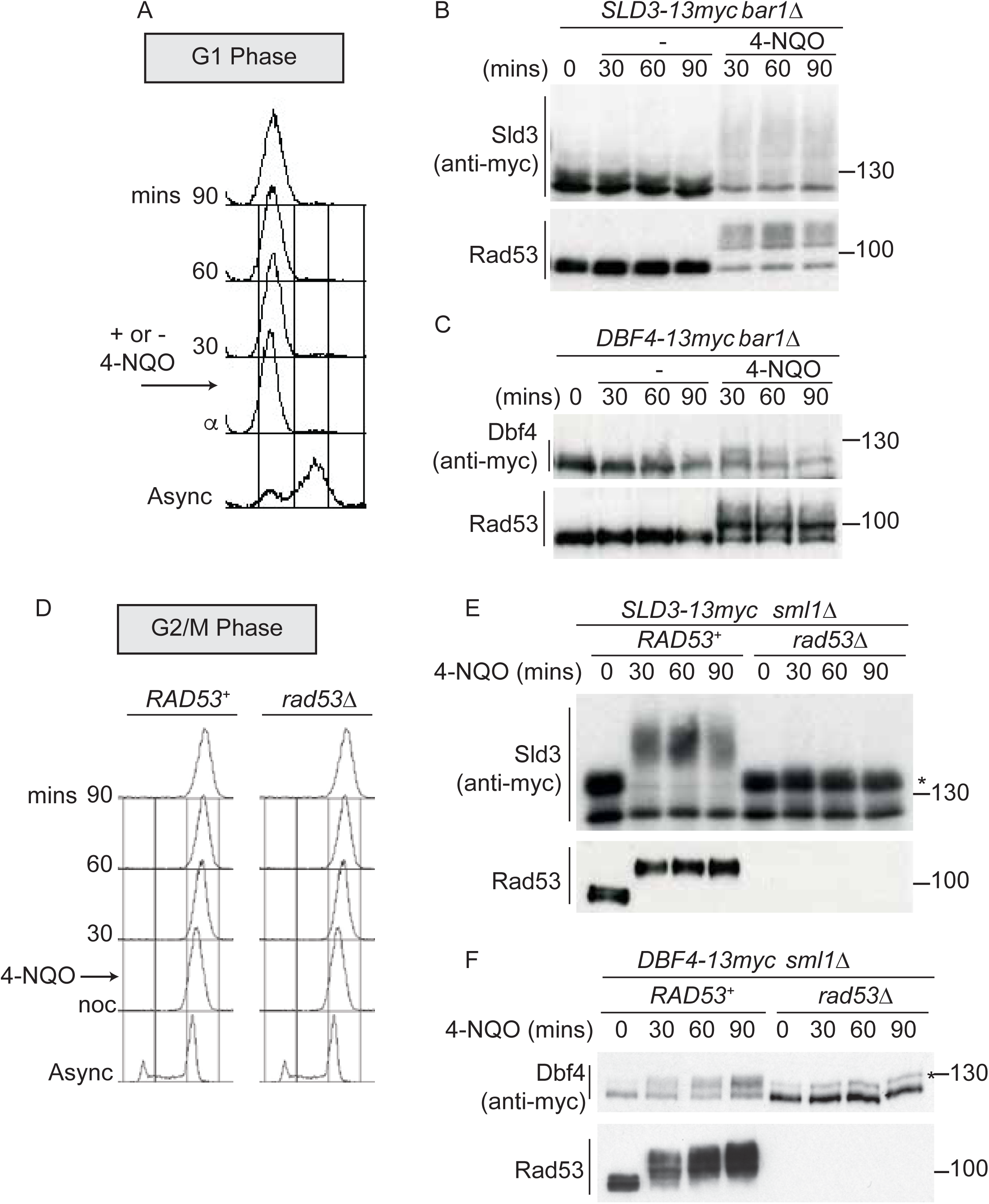
Dbf4 and Sld3 are phosphorylated by Rad53 after DNA damage in G1 and G2 phase. A)Flow cytometry of strains arrested in G1 phase with the mating pheromone alpha factor. Strains were held in G1 phase, with or without the addition of 10µg/ml 4-NQO for the indicated times. All strains are bar1Δ to maintain G1 arrest. B)Western blot of Sld3 (anti-myc) and Rad53 phosphorylation from the experiment outlined in A. Sld3 was resolved on a phos-tag SDS PAGE gel. C)As B, but for Dbf4. Both blots are from SDS-PAGE. D)As A, except strains were arrested in G2/M in nocodazole before the addition of 4-NQO. All strains are sml1Δ. E)Western blot of Sld3 (anti-myc) and Rad53 phosphorylation as in B from the experiment outlined in D. Sld3 was resolved on a phos-tag SDS PAGE gel, * is CDK phosphorylated Sld3. F)As E, but for Dbf4. Dbf4 is phosphorylated by other kinases in G2/M, resulting in residual phosphorylated forms remaining in *rad53Δ* cells *.

To test whether Sld3 and Dbf4 could also be phosphorylated by Rad53 after DNA replication is complete we performed the same experiment as in Figure 1A-C, except in cells arrested in G2/M phase with nocodazole (Figure 1D). Significantly, we observed a Rad53-dependent mobility shift in Sld3 and Dbf4, even in G2/M arrested cells (Figure 1E and 1F). Sld3 and Dbf4 are phosphorylated by other kinases in G2/M, such as by CDK (Holt et al., 2009), giving rise to additional isoforms of Sld3/Dbf4 proteins even in Rad53 null cells (* Figure 1E and 1F). Note that the CDK phosphorylated form of Sld3 is visible in Figure 1E as this is a phos-tag gel.

Previously we have identified the serine and threonine residues in Sld3 and Dbf4 that are directly phosphorylated by Rad53 in S-phase (Zegerman and Diffley, 2010). We mapped 38 such phospho-sites in Sld3 and 19 sites in Dbf4, of which 4 were critical for the Rad53-dependent inhibition of Dbf4. Mutation of these serine/threonine residues to alanine in Sld3 and Dbf4 generated alleles that are refractory to Rad53 phosphorylation in S-phase (Zegerman and Diffley, 2010) and are hereafter referred to as *sld3-A* and *dbf4-A* respectively. We reasoned that if the same sites in Sld3 and Dbf4 are phosphorylated by Rad53 throughout the cell cycle, then *sld3-A* and *dbf4-A* should be defective in Rad53-dependent phosphorylation in G1 and G2/M as well. In G1 phase the Sld3-A protein demonstrated a dramatic loss of Rad53 phosphorylation (Figure 2A), consistent with direct phosphorylation of Sld3 by Rad53 at the same sites as in S-phase. We also observed a similar result with the Dbf4-A protein in G1 phase after Rad53 activation (Figure 2B). As in G1 phase, both Sld3-A and Dbf4-A showed greatly reduced phosphorylation during Rad53 activation in G2/M phase (Figure 2C and 2D). Together, Figures 1 and 2 show that although Sld3 and Dbf4 are considered to be ‘S-phase checkpoint’ substrates of Rad53, they are phosphorylated at the same sites as in S-phase after DNA damage in G1 and G2 phase.

**Figure 2.**
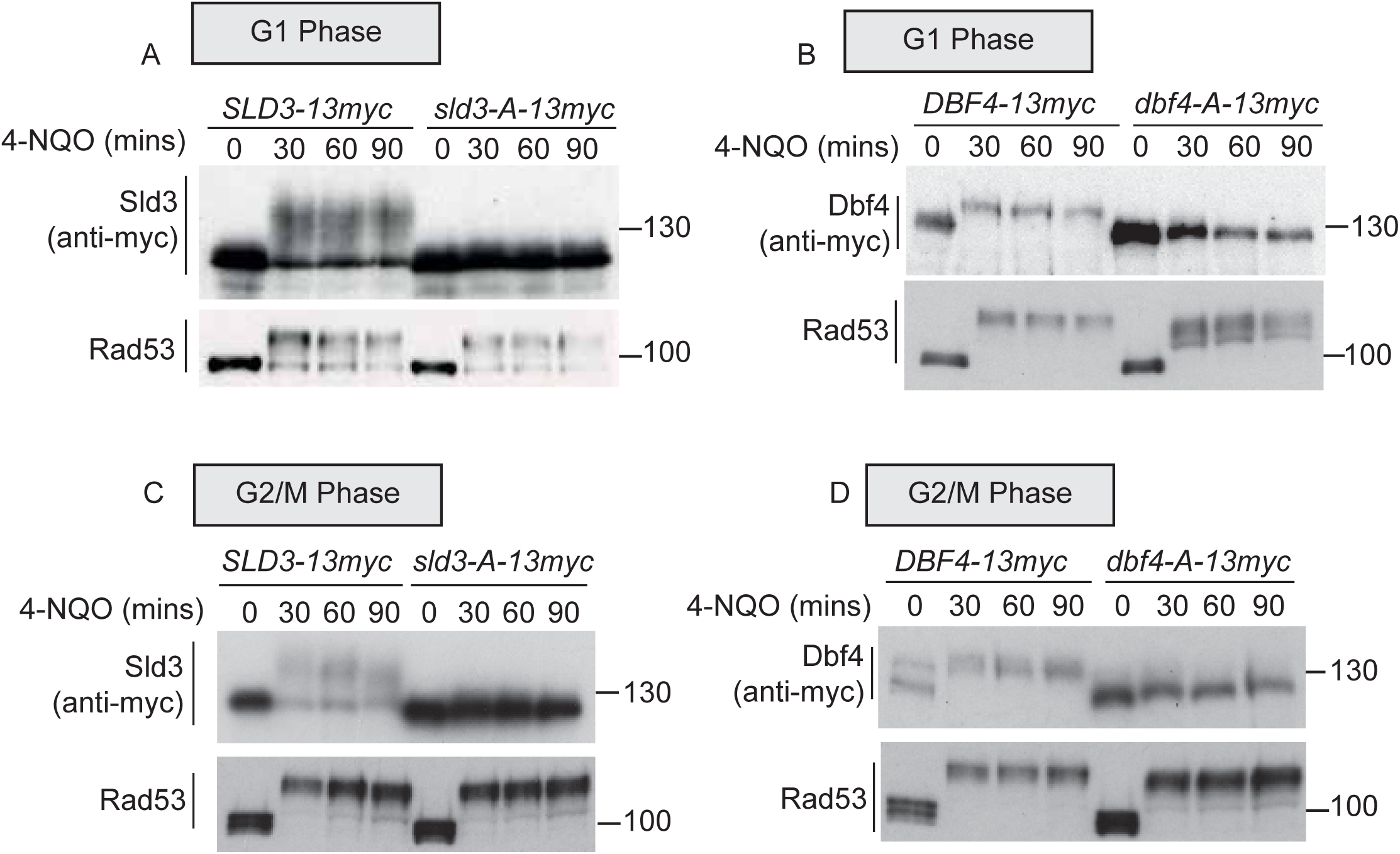
Rad53 phosphorylates Sld3 and Dbf4 in G1 and G2 phase at the same residues as in S-phase. A) and B) as Figure 1 B/C. *sld3-A* and *dbf4-A* refers to mutant alleles with Rad53 phosphorylation sites mutated to alanine (38 sites for Sld3 and 19 sites for Dbf4). All strains are bar1Δ. C)as Figure 1E, except this is Western blot from an SDS-PAGE gel. D)as Figure 1F.

### Rad53-dependent phosphorylation of Sld3 and Dbf4 reduces re-replication in G2 phase

We have previously shown that Rad53 phosphorylates Sld3 and Dbf4 in S-phase to inhibit origin firing (Lopez-Mosqueda et al., 2010; Zegerman and Diffley, 2010). As Sld3 and Dbf4 are phosphorylated at the same sites by Rad53 after DNA damage in both G1 and G2 phase (Figure 1 and 2) we wondered whether this phosphorylation could also be required to inhibit replication initiation outside of S-phase. DNA replication is tightly restricted to S-phase in large part by the action of CDK, which prevents licensing outside of late M/early G1 phase. As a result, transient reduction of CDK-activity in G2/M phase is sufficient to induce re-replication (Dahmann et al., 1995). To test a role for Rad53 phosphorylation of Sld3/Dbf4 in re-replication control we first combined the *sld3-A/dbf4-A* alleles, which cannot be inhibited by Rad53 (Zegerman and Diffley, 2010), with a hypomorphic mutant of the CDK catalytic subunit Cdc28 (*cdc28-as1*). This allele of Cdc28 is analogue sensitive (as) and is inhibited by the addition of the ATP competitive inhibitor 1-NM-PP1. Interestingly we observed that the *sld3-A/dbf4-A* alleles are synthetically sick with *cdc28-as1* in the presence of sub-lethal doses of 1-NM-PP1 (Figure 3A), suggesting that inhibition of Sld3 and Dbf4 by Rad53 is important in cells that have reduced CDK activity.

**Figure 3.**
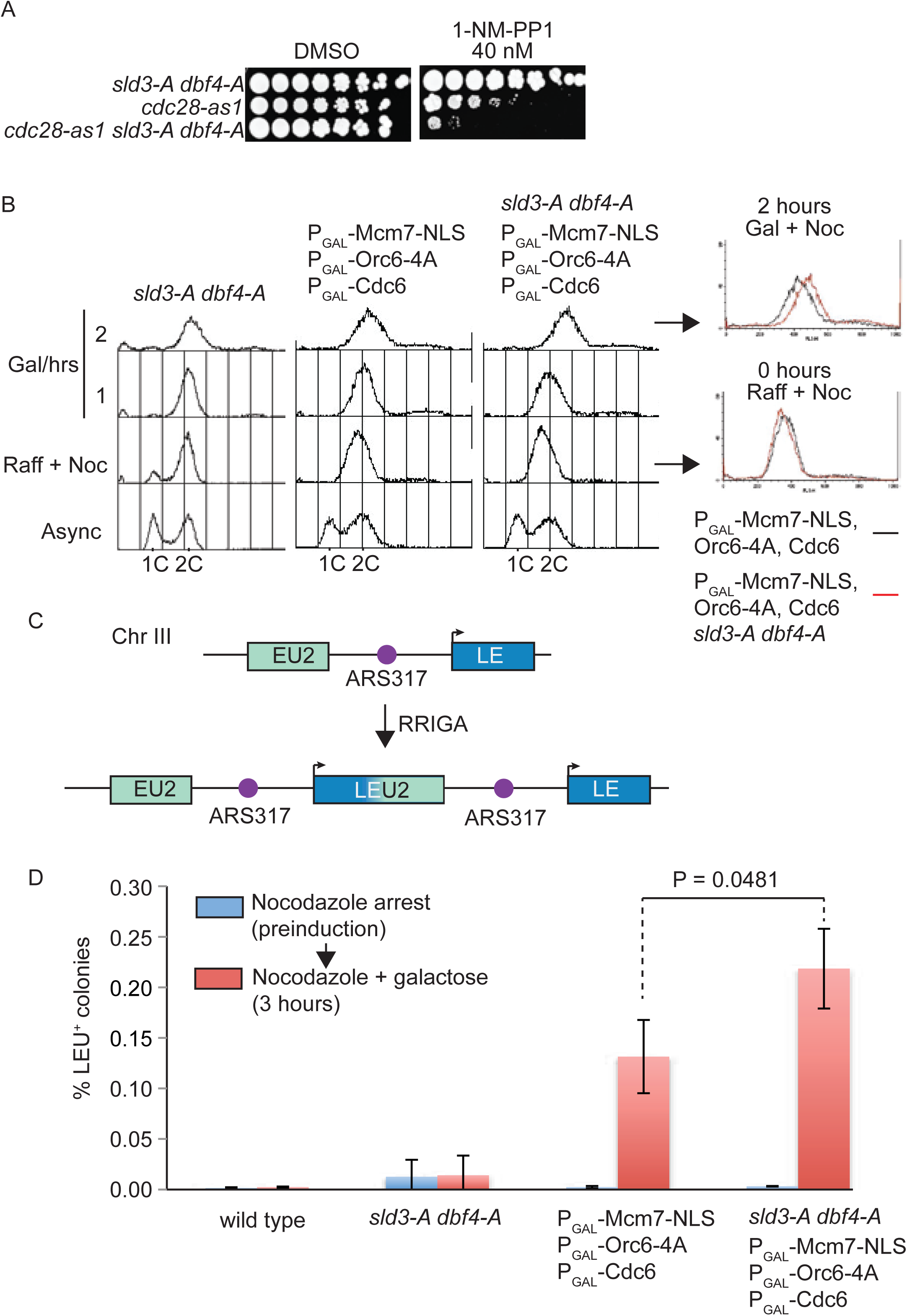
Checkpoint-dependent inhibition of origin firing prevents re-replication in G2 phase. A)Growth assay of the indicated strains B)Flow cytometry of the indicated strains grown overnight in YPraffinose, then arrested in G2/M with nocodazole. After addition of fresh nocodazole, 2% galactose was added for the indicated times to express the licensing mutants. Right, overlay between the 0 and 2 hour timepoints for the licensing mutant strain with (red) or without (black) the *sld3-A/dbf4-A* alleles. C)Schematic diagram of RRIGA assay for gene amplification events. Re-replication of the split LEU2 gene from origin ARS317 results in tandem head to tail gene duplications, leading to a functional LEU2 gene. D)RRIGA assay in C) was performed with the indicated strains. Strains were grown overnight in YPraffinose then arrested in G2/M with nocodazole (pre-induction, blue timepoint). After addition of fresh nocodazole, 2% galactose was added for 3 hours to express the licensing mutants (red timepoint). Cells were plated on YPD (viable cell count) and SC-leu plates (LEU+ count) and the % of LEU+ colonies out of the viable cell population was plotted. N=3, error bars are SD and P value was calculated using an unpaired t-test.

To specifically test whether the Rad53-dependent inhibition of origin firing is important to prevent re-replication, we combined the *sld3-A/dbf4-A* alleles with mutants that circumvent the CDK-dependent inhibition of licensing. Over-expression of Cdc6, forcing the nuclear localisation of the Mcm2-7 complex (through an Mcm7-2xNLS fusion) and mutation of the CDK phosphorylation sites in ORC is sufficient to induce re-replication in G2/M phase (Finn and Li, 2013; Nguyen et al., 2001) and has been shown to induce Rad53 activation (Archambault et al., 2005; Green and Li, 2005). Importantly, conditional over-expression of licensing mutants that cannot be inhibited by CDK combined with *sld3-A* and *dbf4-A* led to an increase in the total re-replication in nocodazole arrested cells (Figure 3B – compare FACS overlay red vs black). This suggests that Rad53-dependent inhibition of replication initiation can reduce inappropriate replication in G2 phase.

One of the consequences of re-replication is the generation of head-to-tail tandem gene amplifications, a process termed RRIGA (re-replication induced gene amplification, Green et al., 2010). To examine whether the Rad53-dependent inhibition of origin firing helps to prevent RRIGA we adapted an assay to quantitatively assess gene amplification events in G2/M arrested cells (Finn and Li, 2013). Briefly a marker gene (in this case LEU2, which allows growth on media lacking leucine) was split with some remaining homology across an origin that re-initiates when licensing control is lost (ARS317). Re-initiation at ARS317 followed by fork-breakage and strand annealing at the regions of LEU2 homology results in gene amplification and the generation of a functional LEU2 gene (Figure 3C and Supplementary Figure 2). In this assay, as in Figure 3B, the re-replication mutants were induced only in G2/M arrested cells. In contrast to wild type yeast or the *sld3-A/dbf4-A* strain alone, expression of the licensing mutants by the addition of galactose resulted in a large increase in RRIGA events, as expected (Figure 3D). Importantly RRIGA events were even greater when the mutants that allow licensing in the presence of CDK were combined with the *sld3-A*/*dbf4-A* alleles (Figure 3D). This assay demonstrates that the checkpoint kinase Rad53 indeed reduces gene amplification events after re-replication through inhibition of Sld3 and Dbf4, even in G2/M arrested cells.

### Rad53 prevents precocious origin firing after DNA damage in G1 phase

DNA damage in G1 phase delays the G1/S transition, which from humans to yeast, involves the checkpoint kinase-dependent down-regulation of G1/S cyclins, delaying cell cycle entry (Bertoli et al., 2013; Lanz et al., 2019; Shaltiel et al., 2015; Sidorova and Breeden, 1997). Here we have shown that DNA damage in G1 phase also results in the inhibitory checkpoint phosphorylation of two replication initiation factors, Sld3 and Dbf4 (Figure 1 and 2), suggesting that this might be an additional mechanism to prevent premature DNA replication at the G1/S transition (Figure 4A). To specifically analyse the consequences of DNA damage in G1 phase we added 4-NQO to G1 arrested yeast cells and then released cells into S-phase in fresh medium without 4-NQO. Crucially, this approach resulted in robust Rad53 activation in G1 phase, such that cells enter S-phase with an already active checkpoint (Figure 4D and Supplementary Figure 3C). Rad53 activation in G1 phase resulted in the slowing of the G1/S transition as detected by the delay in budding (a G1 cyclin mediated event) and delay in DNA synthesis (Supplementary Figure 3A-B). Despite this the *sld3-A dbf4-A* alleles caused little difference in S-phase progression after DNA damage in G1 phase, compared to the wild type strain (Supplementary Figure 3A). As Rad53 is known to inhibit CDK activation through phosphorylation of the Swi6 subunit of the transcriptional activator SBF (Sidorova and Breeden, 1997), we wondered whether Rad53-dependent inhibition of both origin firing and G1/S transcription might prevent precocious DNA replication after damage in G1 phase (Figure 4A). To test this we over-expressed a truncated form of the SBF transcription factor Swi4 (Swi4-t), which lacks the C-terminus required for interaction with Swi6 and thus cannot be inhibited by Rad53 (Sidorova and Breeden, 1997). Over-expression of Swi4-t indeed resulted in faster progression through the G1/S transition in the presence of 4-NQO (Supplementary Figure 3D), as expected (Sidorova and Breeden, 1997). Importantly the combination of expression of Swi4-t together with *sld3-A dbf4-A* resulted in much faster S-phase progression after DNA damage in G1 phase compared to Swi4-t expression alone (Figure 4B). These differences in the onset of DNA replication between *swi4-t* with and without *sld3-A/dbf4-A* were not due to differences in the G1/S transition as these strains budded at the same time (Figure 4C) and both strains also exhibited similar levels of Rad53 activation (Figure 4D). Together this suggests that Rad53 activation in G1 phase prevents precocious DNA replication initiation by not only inhibiting G1/S transcription, but also by inhibiting Sld3 and Dbf4.

**Figure 4.**
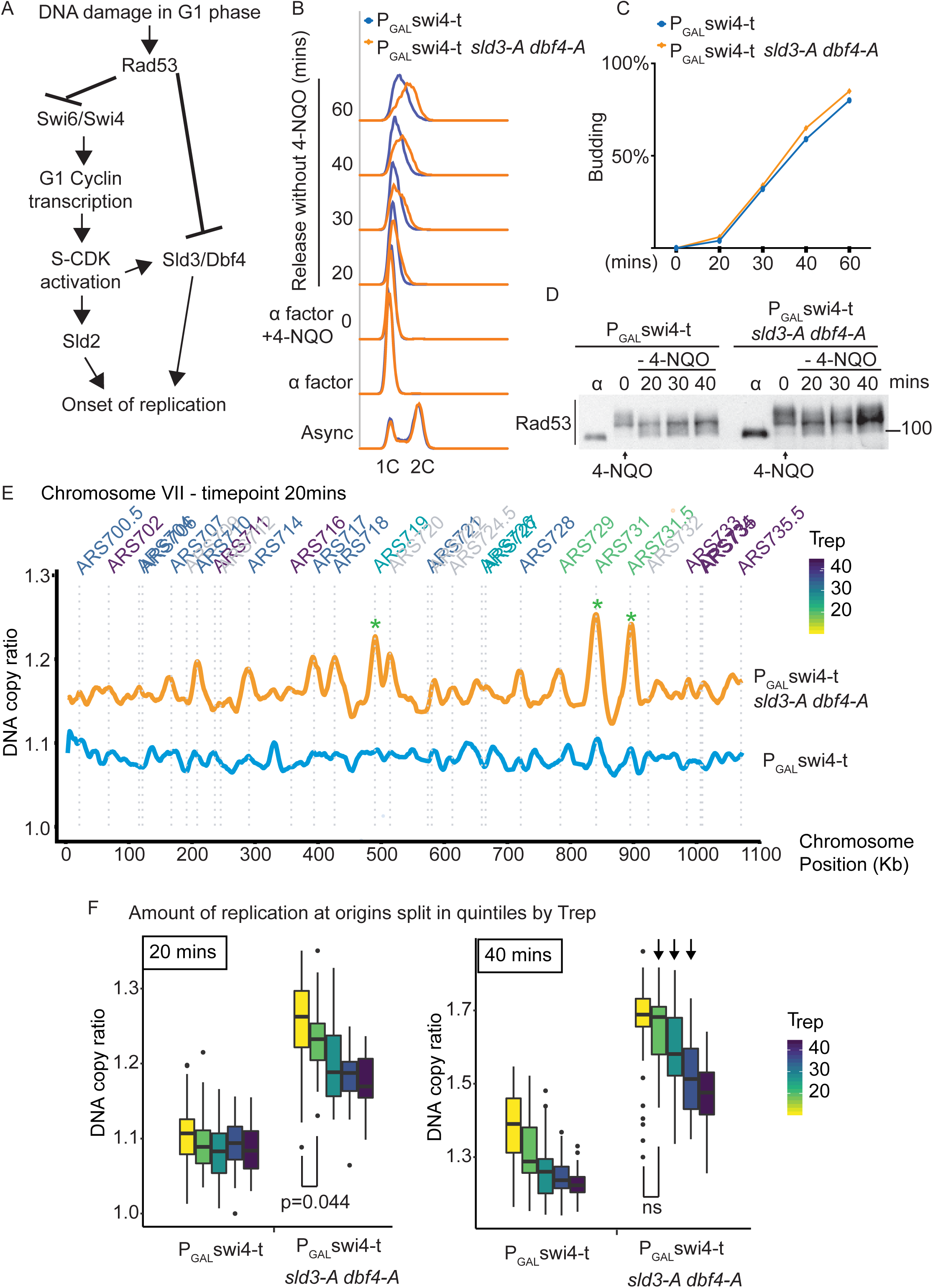
Checkpoint-dependent inhibition of origin firing prevents premature replication initiation at all origins at the G1-S transition. A)Activation of Rad53 in G1 phase can delay genome duplication by at least 2 mechanisms; by inhibition of the transcriptional activator Swi6, which is required for G1- and subsequently S-phase cyclin transcription and by inhibition of the origin firing factors Sld3 and Dbf4. B)Flow cytometry of the indicated strains grown overnight in YPraffinose, then arrested in G1 phase with alpha factor. Cells were held in fresh alpha factor, while 2% galactose and 0.5μg/ml 4-NQO was added for 30 minutes (0 timepoint) before washing and release from alpha factor arrest into fresh YPgal medium without 4-NQO. C)Budding index from the experiment in B. Timepoint 0 refers to cells held in alpha factor + galactose + 0.5μg/ml 4-NQO for 30mins. D) Rad53 Western blot from experiment in B. E)Copy number analysis of chromosome VII of the indicated strains 20 mins after release as in B-D. The y-axis ratio refers to the amount of DNA at the 20 mins timepoint divided by the DNA copy number in G1 phase. Known origins are annotated above the replication profile and coloured according to their normal median replication time (T_rep_). F)Box plots of the amount of replication at all origins, split into equal quintiles depending on their normal median firing time (T_rep_). Arrows in time-point 40 mins indicate that later firing origins also initiate by 40mins in the *swi4-t sld3-A dbf4-A* strain. For example the yellow and green quintiles are significantly different at 20mins, but non-significantly different (ns) at 40mins. P-values are from t-tests.

Activation of Rad53 during S-phase caused by fork stalling/DNA damage at early replicons results in inhibition of subsequent (late) origin firing, which in yeast is mediated by inhibition of Sld3 and Dbf4 (Lopez-Mosqueda et al., 2010; Zegerman and Diffley, 2010). We therefore wondered whether the accelerated S-phase we observe when we combine *swi4-t* with *sld3-A dbf4-A*, is simply due to the canonical S-phase checkpoint inhibition of late origin firing or whether by activating Rad53 in G1 phase we are actually causing a delay in genome duplication from all origins. To assess this we analysed the replication dynamics of the time-course in Figure 4B-D by high-throughput sequencing and copy number analysis. At the earliest time-point (20mins), while the *swi4-t* over-expressing strain alone had barely begun to replicate (Figure 4E, chromosome VII as an example), the *swi4-t sld3-A dbf4-A* strain showed peaks of replication initiation at the earliest firing origins (for example see *, Figure 4E), even though Rad53 is highly activated (Figure 4D). By analysis of initiation at all origins, split into quintiles according to their normal firing time, we observe much greater firing of early origins in the *swi4-t sld3-A dbf4-A* strain compared to swi4-t alone throughout the time-course (Figure 4F and Supplementary Figure 4). Over time, we also observe an increase in later firing origins in the *swi4-t sld3-A dbf4-A* strain (arrows Figure 4F and Supplementary Figure 4), suggesting that the relative timing of origin firing is not affected. Together this demonstrates that activation of Rad53 and inhibition of Sld3 and Dbf4 in G1 phase contributes to the mechanism preventing the onset of DNA replication from all origins, not just late firing origins, in the presence of DNA damage (Figure 4A).

If the checkpoint-mediated inhibition of G1/S transcription and Sld3/Dbf4 both contribute to prevent precocious S-phase entry then we hypothesised that loss of both pathways should show synthetic lethality in the presence of DNA damage. We have previously conducted genetic interaction analysis of the *sld3-A/dbf4-A* alleles with the yeast whole genome gene knock-out collection in the presence of the DNA damaging agent phleomycin (Morafraile et al., 2019). Significantly loss of function of genes that result in a delay in the G1/S transition, such as *CLN2, SWI4* and *BCK2* (Di Como et al., 1995) improved the growth of *sld3-A/dbf4-A* in the presence of phleomycin (suppressors, Supplementary Figure 5), whereas loss of function of genes that would result in the acceleration of G1/S, such as *WHI5* and *SIC1* (Bertoli et al., 2013) were synthetic sick with the *sld3-A/dbf4-A* alleles (enhancers, Supplementary Figure 5). These genetic interactions are consistent with an important role for Rad53-dependent inhibition of origin firing in preventing precocious replication after DNA damage in G1 phase.

Here we show that the two critical targets of the S-phase checkpoint mediated inhibition of origin firing, Sld3 and Dbf4, are actually regulated by Rad53 after DNA damage throughout the cell cycle (Figures 1 and 2). This has important implications for understanding the consequences of inappropriate re-replication in human cells, where the role of the checkpoint differs depending on the cell cycle phase in which re-replication occurs (Klotz-Noack et al., 2012; Liu et al., 2007). By combining tight cell cycle arrests with the separation-of-function mutants, *sld3-A dbf4-A*, we show specifically that the checkpoint-dependent inhibition of origin firing limits further re-replication and gene amplifications in G2 phase when licensing control is compromised (Figure 3). This pathway is likely to be evolutionarily conserved, as checkpoint activation in S-phase in human cells also appears to limit re-replication through inhibition of Dbf4 (Lee et al., 2012). As tandem head-to-tail duplications are a prominent feature of many cancers (Menghi et al., 2018), knowledge of the pathways that prevent this form of structural variation may be important for understanding oncogenesis.

In addition to preventing re-replication in G2, we show that the checkpoint also inhibits the replication initiation factors Sld3 and Dbf4 to delay origin firing after DNA damage in G1 phase (Figure 4). This likely increases the time for DNA repair to occur before replication begins and may also serve to increase the window of time where origin firing and licensing are mutually exclusive, preventing re-replication. It is interesting that failure to inhibit Sld3 and Dbf4 alone has little effect on the G1/S transition (Supplementary Figure 3A), probably because Sld3 (and Sld2) act downstream of CDK activation (Figure 4A). Inhibition of origin firing may therefore be a failsafe mechanism when inhibition of G1/S CDK activity is incomplete. Mutations in genes such as Rb and p53 that control the G1/S transition and the G1 checkpoint response respectively are amongst the most common mutations in cancers (Malumbres and Barbacid, 2001; Massague, 2004). Work from yeast to humans has shown that defects in the G1/S transition results in increased dependence on the checkpoint kinases for survival (Rundle et al., 2017; Sidorova and Breeden, 2002). The checkpoint inhibition of all origin firing as a failsafe to prevent precocious S-phase entry described in this study (Figure 4) may provide a potential mechanistic rationale for the selective targeting of p53/Rb mutant cancers using Chk1 and ATR inhibitors, which are currently in clinical trials (Bradbury et al., 2020).

## Methods

### Strains and Growth Conditions

Cell growth, arrests, flow cytometry and yeast protein extracts were as previously described (Zegerman and Diffley, 2010). All the strains used in this work are derived from W303 (*ade2-1 ura3-1 his3-11,15 trp1-1 leu2-3,112 can1-100, rad5-535)*

| Strain | Genotype | Source |
| --- | --- | --- |
| yPZ 1223 | *MATa Sld3-10his13myc::KanMX bar1Δ::hisG* | this work |
| yPZ 1319 | *MATa sld3-38A-10his13myc::KanMX bar1Δ::hisG* | this work |
| yPZ 1317 | *MATa Dbf4-13myc::KanMX bar1Δ::hisG* | this work |
| yPZ 4018 | *MATa YCLWdelta5::HphMX-EU2-ARS317-LEU::NFS1*<br>*Mcm7-2xNLS::AbA ura3::P<sub>GAL</sub>-Cdc6-13myc::URA3 orc6-4A* | this work |
| yPZ 705 | *MATa* | PZ lab |
| yPZ 4076 | *MATa trp1::P<sub>GAL</sub>-swi4-t::TRP1* | this work |
| yPZ 4137 | *MATa dbf4-4A::HIS3 sld3-38A-10his13myc::KanMX*<br>*trp1::P<sub>GAL</sub>-swi4-t::TRP1* | this work |
| yPZ 917 | *MATa dbf4-4A::HIS3 sld3-38A-10his13myc::KanMX* | PZ lab |
| yPZ 1523 | *MATa sml1Δ::URA3 rad53Δ::LEU2 Sld3-10his13myc::KanMX*<br>*bar1Δ::hisG* | this work |
| yPZ 1522 | *MATa sml1Δ::URA3 Sld3-10his13myc::KanMX bar1Δ::hisG* | this work |
| yPZ 1198 | *MATa cdc28-as1 (F88G)* | this work |
| yPZ 1767 | *MATa cdc28-as1 (F88G) dbf4-4A::HIS3*<br>*sld3-38A-10his13myc::KanMX* | this work |
| yPZ 4085 | *MATa YCLWdelta5::HphMX-EU2-ARS317-LEU::NFS1*<br>*trp1::orc6-4A-P<sub>GAL1-10</sub>-Mcm7-2xNLS::TRP1*<br>*ura3::P<sub>GAL</sub>-Cdc6-13myc::URA3* | this work |
| yPZ 4089 | *YCLWdelta5::HphMX-EU2-ARS317-LEU::NFS1*<br>*trp1::orc6-4A-P<sub>GAL1-10</sub>-Mcm7-2xNLS::TRP1*<br>*sld3-38A-10his13myc::KanMX dbf4-4A::HIS3*<br>*ura3::P<sub>GAL</sub>-Cdc6-13myc::URA3* | this work |
| yPZ 125 | *MATa Dbf4-13myc::KanMX* | PZ lab |
| yPZ 170 | *MATa dbf4Δ::TRP1 his3::P<sub>DBF4</sub>-dbf4-19A-13myc::KanMX::HIS3* | PZ lab |
| yPZ 2 | *MATa sld3-38A-10his13myc::KanMX* | PZ lab |
| yPZ 52 | *MATa Sld3-10his13myc::KanMX* | PZ lab |
| yPZ 1471 | *MATa sml1Δ::URA3 rad53Δ::HphNT1 Dbf4-13myc::KanMX*<br>*bar1Δ::hisG* | this work |
| yPZ 1473 | *MATa sml1Δ::URA3 Dbf4-13myc::KanMX bar1Δ::hisG* | this work |
| yPZ 520 | *MATa sml1Δ::URA3 Sld3-13myc::KanMX* | PZ lab |
| yPZ 89 | *MATa sml1Δ::URA3 rad53Δ::LEU2 Sld3-13myc::KanMX* | PZ lab |
| yPZ 519 | *MATa sml1Δ::URA3 Dbf4-13myc::KanMX* | PZ lab |
| yPZ 228 | *MATa sml1Δ::URA3 rad53Δ::LEU2 Dbf4-13myc::KanMX* | PZ lab |

*dbf4-4A* refers to the rad53 site mutant. It has serine/threonine to alanine mutations at amino acids: 518, 521, 526, 528.

*dbf4-19A* refers to the rad53 site mutant. It has serine/threonine to alanine mutations at amino acids 53, 59, 188, 192, 203, 222, 224, 226, 228, 318, 319, 328, 374, 375, 377, 518, 521, 526, 528.

*sld3-38A* refers to the rad53 site mutant. It contains serine/threonine to alanine mutations at amino acids 306, 310, 421, 434, 435, 438, 442, 445, 450, 451, 452, 456, 458, 459, 479, 482, 507, 509, 514, 519,521,524, 540, 541, 546, 547, 548, 550, 556, 558, 559, 565, 569, 582, 607, 653 and 654. 539 is mutated to arginine.

*Orc6-4A* refers to CDK sites 106, 116, 123 and 146 mutated to alanine.

### Western blot

Western blots were performed as previously described (Can et al., 2019). Rad53 was detected with ab104232 (Abcam, dilution 1:5000).

### Replication profiles

Yeast genomic DNA was extracted using the smash & grab method (https://fangman-brewer.genetics.washington.edu/smash-n-grab.html). DNA was sonicated using the Bioruptor Pico sonicator (Diagenode) and the libraries were prepared according to the TruSeq Nano sample preparation guide from Illumina. To generate replication timing profiles, the ratio of uniquely mapped reads in the replicating samples to the non-replicating samples was calculated following (Batrakou et al., 2020). Replication profiles were generated using custom R scripts and smoothed using a moving average. The values of Trep were taken from OriDB (Siow et al., 2012).

### RRIGA assay using split LEU2 marker

Cells were pre-grown under permissive conditions in YPraff overnight 30·C. At 1×10^7^ cells/ml, nocodazole (2mg/ml in DMSO) was added to a final concentration of 10µg/ml. Cells were arrested for 90mins at 30·C (uninduced timepoint) and then galactose was added to a final concentration of 2% + fresh nocodazole for 3 hours (induced timepoint).

For each timepoint a 0.5ml sample was spun at 3.2K for 1min in benchtop centrifuge, cells were washed with 1ml sterile water to remove YPD, respun and resuspended in 0.5ml sterile water. Cells were sonicated briefly to ensure cells are separated then a serial dilution was made into sterile water as follows:

dilution 1: 10µl cells + 990µl water = approx 1×10^5^ cells/ml = 1×10^2^ cells/µl dilution 2: 10µl dilution 1 + 990µl water = approx 1×10^3^ cells/ml = 1 cell/µl

100µl of dilution 2 was plated on YPD plates in triplicate for the viability calculation. 100µl and 10µl of undiluted cells were plated in triplicate on SC-leu plates to obtain the fraction of viable cells that are LEU+ before and after induction of re-replication. Plates were incubated at 25·C for 48hrs before colonies were counted. To calculate the percentage of cells in the population that were LEU+ the number of LEU+ colonies per ml was divided by the number of viable cells per ml.

## Acknowledgments

We thank members of the Zegerman lab for critical reading of the manuscript. Work in the PZ lab was supported by AICR 10-0908, Wellcome Trust 107056/Z/15/Z, Cancer Research UK C15873/A12700 and Gurdon Institute funding (Cancer Research UK C6946/A14492, Wellcome Trust 092096). Part III undergraduate student DA was supported by the Department of Biochemistry. MS was funded by the BBSRC BB/M011194/1. GC was supported by a Turkish government grant and a Raymond and Beverley Sackler studentship.

## Author Contribution

All authors performed and designed the experiments. PZ wrote the paper.

## Supplementary Figure legends

**Supplementary Figure 1.**
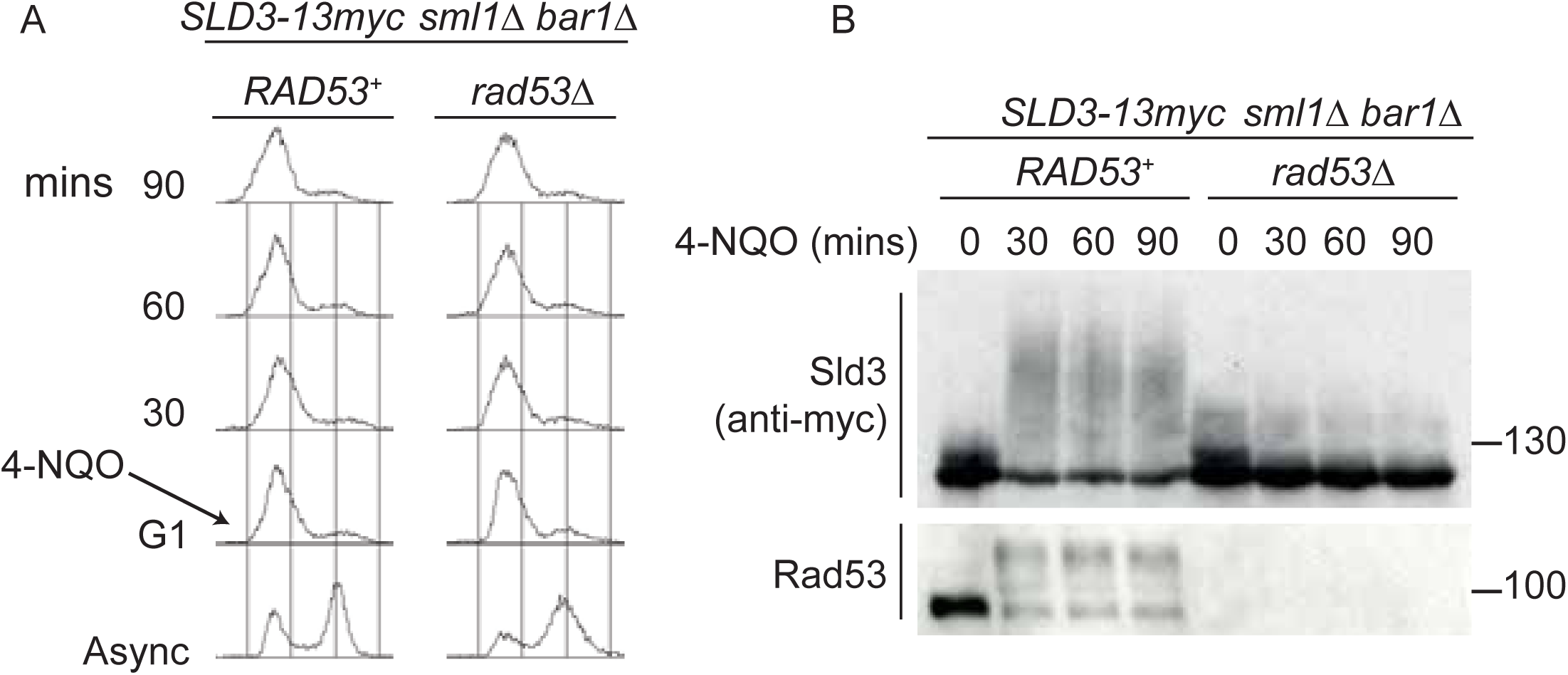
Sld3 phosphorylation in G1 phase after DNA damage is Rad53-dependent. A)Flow cytometry of strains arrested in G1 phase with the mating pheromone alpha factor. Strains were held in G1 phase, with the addition of 10µg/ml 4-NQO for the indicated times. All strains are *bar1Δ* to maintain G1 arrest and also *sml1Δ*. B)Western blot of Sld3 and Rad53 phosphorylation from the experiment outlined in A. Sld3 was resolved on a phos-tag SDS PAGE gel.

**Supplementary Figure 2.**
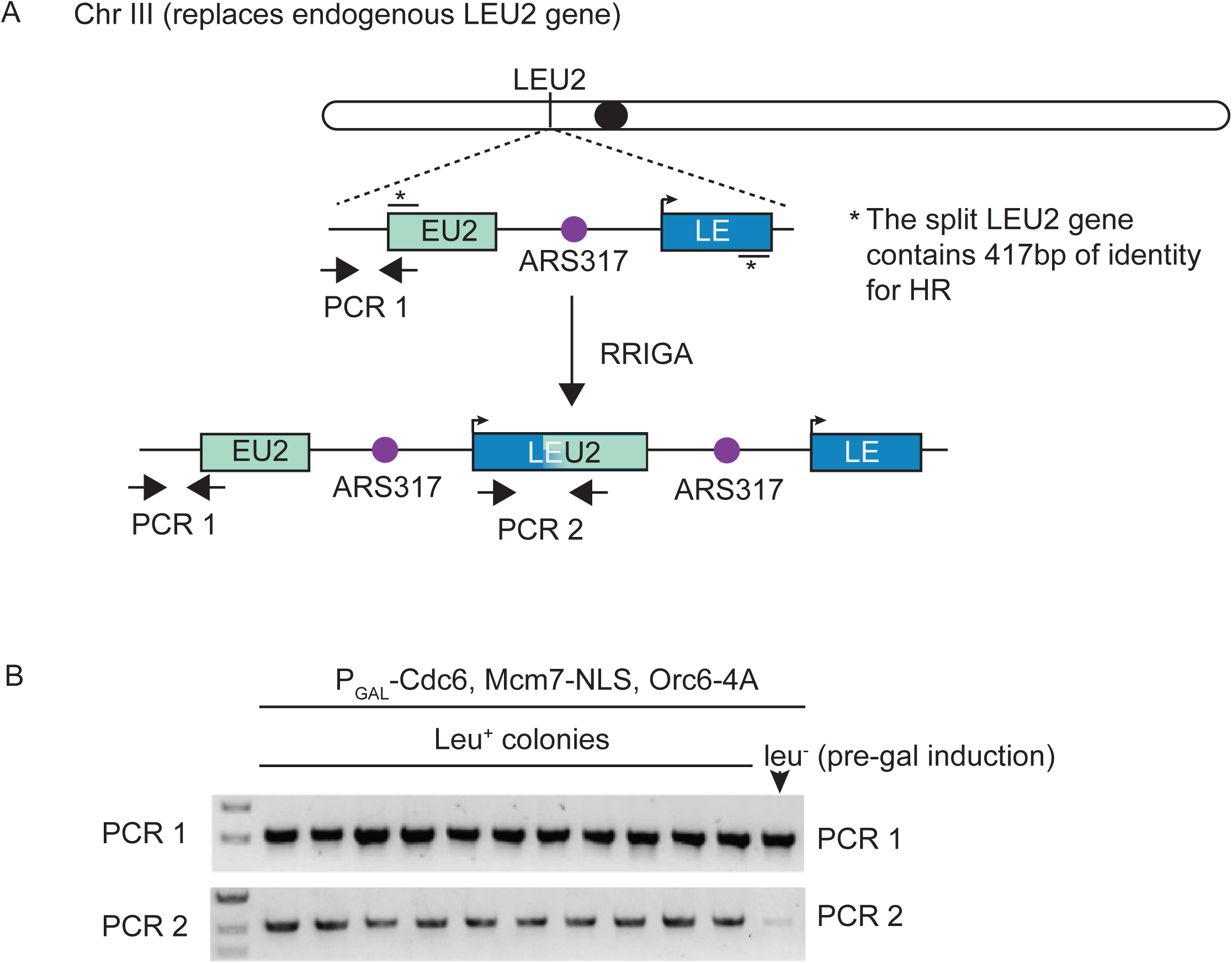
RRIGA assay using split LEU2 marker. A. Schematic diagram of the RRIGA assay. Endogenous LEU2 was replaced with a split LEU2 marker, separated by the re-replication origin ARS317. The two non-functional halves contain 417bp of identity *. Re-initiation at ARS317 followed by fork-breakage and strand annealing at the regions of LEU2 homology results in gene amplification and the generation of a functional LEU2 gene. B. Verification of RRIGA by PCR using the amplicons as depicted in A. The parental strain (which is leu-) was induced to re-replicate in the presence of galactose and cells were plated on SC-leu plates. 10 independent colonies were assayed for the presence of the duplication and all were positive.

**Supplementary Figure 3.**
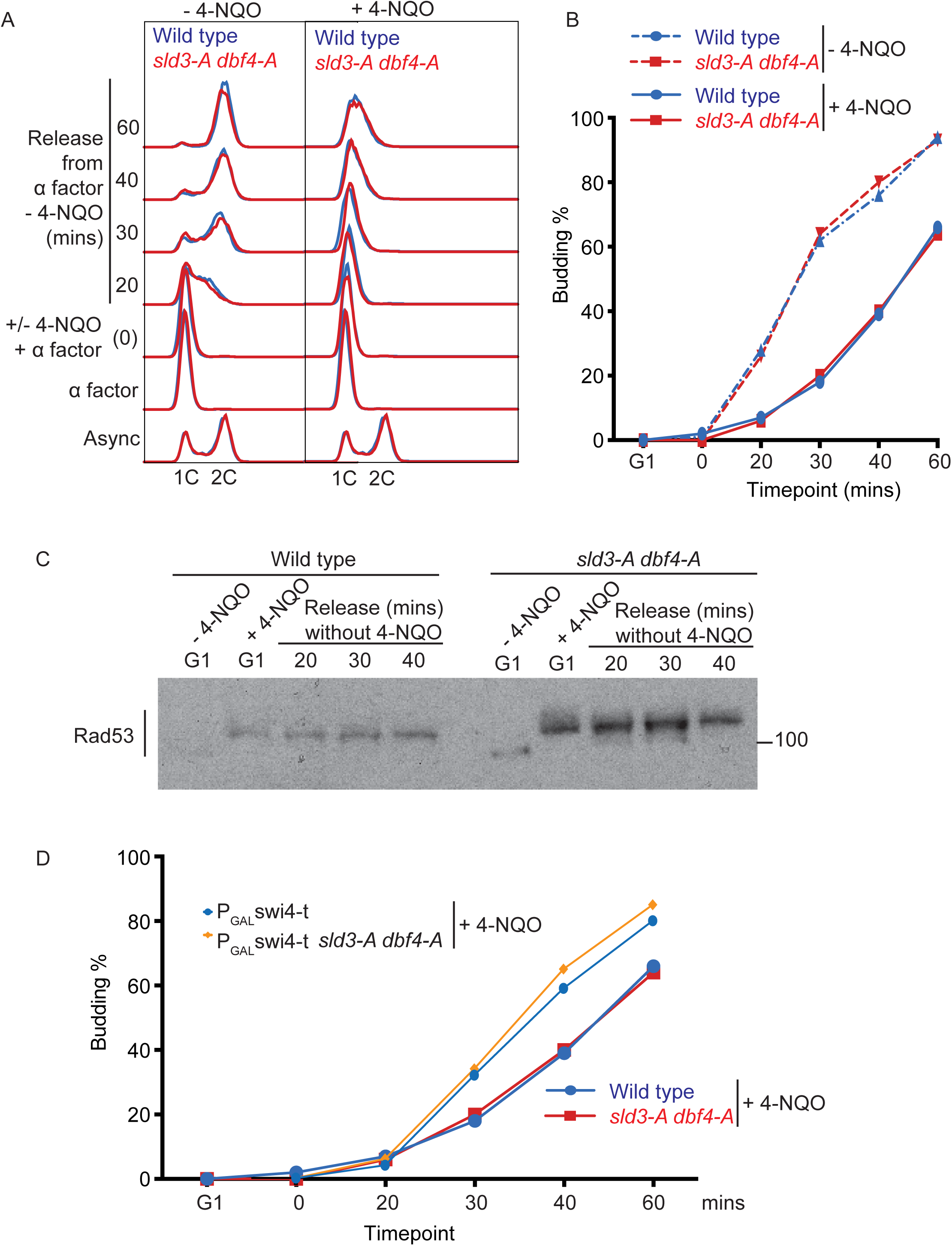
4-NQO addition in G1 phase delays the G1/S transition. A. Flow cytometry of the indicated strains, which were grown overnight in YPraffinose and arrested in G1 phase with alpha factor (G1 time point). Strains were then held in G1 phase for an extra 30 minutes by the addition of fresh alpha factor plus galactose, with or without 0.5μg/ml 4-NQO. Cells were washed into fresh YPgalactose medium to release into S-phase. Note that YPraffinose and galactose medium was used in this experiment to match the exact conditions that were used in the main Figure 4B with the *P*_*GAL*_*-swi4-t* strains, allowing a direct comparison, as in D. B. Budding index (a G1 cyclin mediated event) from experiment in A. C. Rad53 western blot from the 4-NQO treated samples in A. D. Overlay of budding profiles (from Figure 4C and this Figure B) between strains expressing or not expressing Swi4-t. Swi4-t, which cannot be inhibited by Rad53, causes earlier activation of G1 cyclin/CDK as shown here by budding.

**Supplementary Figure 4.**
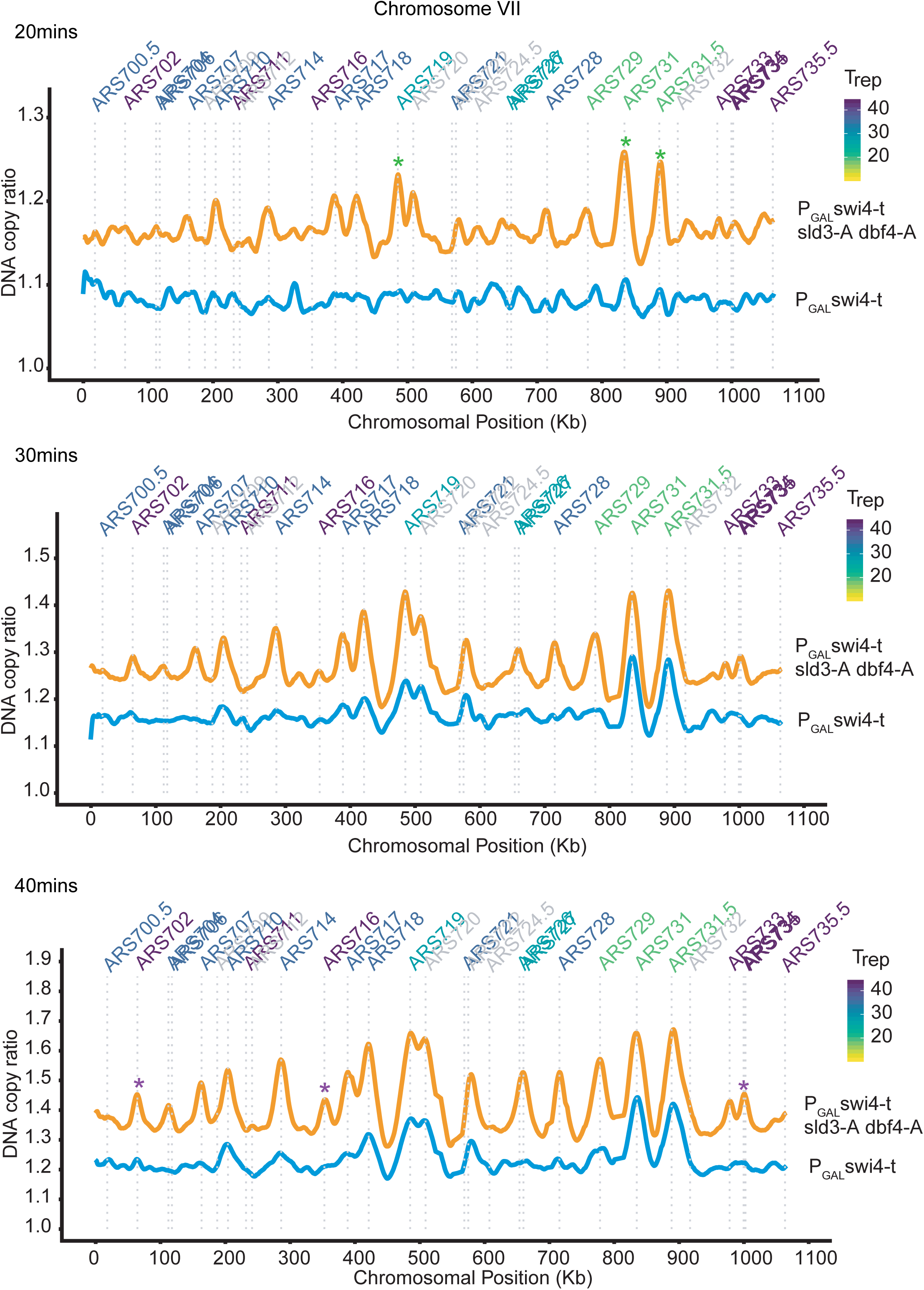
Rad53 activation in G1 phase inhibits all origin firing through phosphorylation of Sld3 and Dbf4. Replication profile of Chromosome VII as an example at 20, 30 and 40mins from the experiment in Figure 4B. Green * are examples of very early firing origins, which initiate at the 20min timepoint in the *P*_*GAL*_*-swi4-t sld3-A dbf4-A* strain. Purple * are examples of late firing origins which do not initiate at the 20min timepoint, but are clearly firing by the 40 min timepoint in the *P*_*GAL*_*-swi4-t sld3-A dbf4-A* strain. This shows that the relative timing of origin firing is not affected in the *P*_*GAL*_*-swi4-t sld3-A dbf4-A* strain.

**Supplementary Figure 5.**
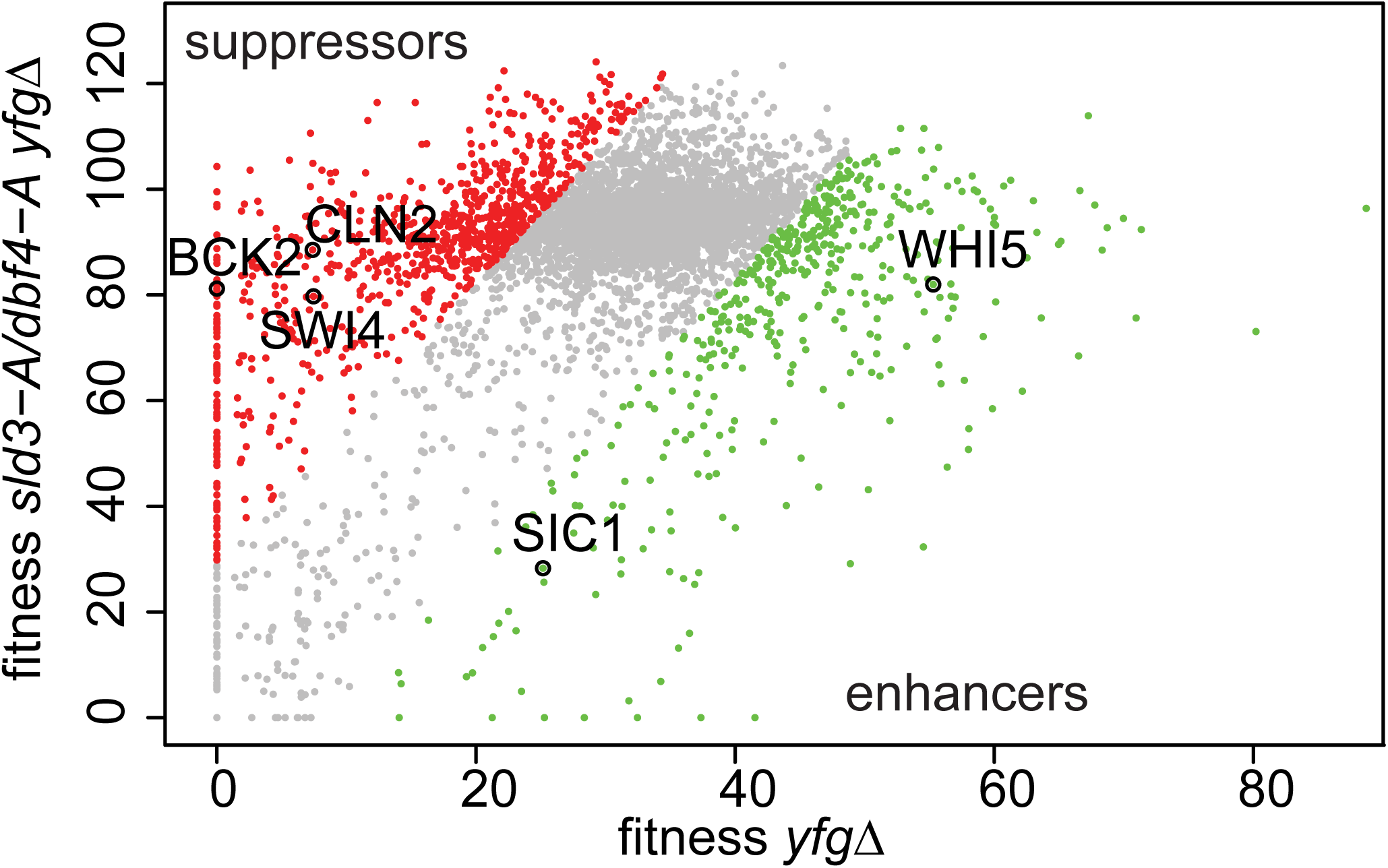
Genetic interactions of the *sld3-A dbf4-A* alleles with mutants that either accelerate or delay the G1/S transition. Scatter plot of the fitness of the yeast genome knock out collection grown in 0.5μg/ml phleomycin with (y-axis) or without (x-axis) the *sld3-A dbf4-A* alleles. Each dot corresponds to a different gene deletion. The top 25% of gene deletions (yfg = your favourite gene) that significantly enhance (green) or suppress (red) the fitness of *sld3-A dbf4-A* are indicated. This data was originally published in (Morafraile et al., 2019).

